# Decreased voluntary alcohol intake and ventral striatal epigenetic and transcriptional remodeling in male Acss2 KO mice

**DOI:** 10.1101/2023.12.04.569964

**Authors:** Gabor Egervari, Greg Donahue, Natalia Quijano-Carde, Connor Hogan, Erica M Periandri, Vanessa Fleites, Desi C Alexander, Mariella De Biasi, Shelley L Berger

**Author notes:** Corresponding author. (G.E.); (S.L.B.).

## Abstract

Metabolic-epigenetic interactions are emerging as key pathways in regulating alcohol-related transcriptional changes in the brain. We previously demonstrated that this is mediated by the metabolic enzyme Acetyl-CoA synthetase 2 (Acss2), which is nuclear and chromatin-bound in neurons. Mice lacking Acss2 fail to deposit alcohol-derived acetate onto histones in the brain and show no conditioned place preference for ethanol reward. Here, we explored the role of this pathway during voluntary alcohol intake. We found that Acss2 KO mice consumed significantly less alcohol during drinking-in-the-dark, and this effect was primarily driven by males. We performed genome-wide transcriptional profiling of 7 key brain regions implicated in alcohol and drug use, and found that, following drinking, Acss2 KO mice exhibited blunted gene expression in the ventral striatum, and similar to the behavioral differences, transcriptional dysregulation was more pronounced in male mice. Further, we found that the gene expression changes were associated with depletion of ventral striatal histone acetylation (H3K27ac) in Acss2 KO mice compared to WT. Taken together, our data suggest that Acss2 plays an important role in orchestrating ventral striatal epigenetic and transcriptional changes during voluntary alcohol drinking, especially in males. Consequently, targeting this pathway could be a promising new therapeutic avenue.

## INTRODUCTION

Alcohol use disorder (AUD) and other abused substances impose a tremendous burden on society as conventional pharmacotherapies lack substantial and durable efficacy^1-3^. Epigenetic mechanisms^4^, such as histone acetylation, which dynamically regulate gene expression are strongly linked both to acute alcohol exposure and to AUD^5-12^. Therefore, emerging new approaches based on epigenetic mechanisms hold considerable promise for AUD and psychiatric illness in general^13-21^. For example, histone deacetylase inhibitors were proposed to alleviate depression-like symptoms during ethanol-withdrawal by normalizing the epigenetic landscape of the brain^16,17^.

Epigenetic mechanisms are strongly influenced by metabolic processes^22,23^. In fact, metabolism of ethanol itself has been proposed to contribute to alcohol-induced epigenetic alterations^8^. Until recently, this has been mostly attributed to changes in available substrates and cofactors^8,24,25^, increases in the NADH/NAD ratio^8,26^, or reactive oxygen species production^27^. Importantly, however, alcohol metabolism gives rise to surges of acetate in the systemic circulation following alcohol exposure^28^, which can directly affect brain function^29^.

Recently, we described a novel, direct and rapid mechanism for alcohol-driven brain histone acetylation governed by a metabolic enzyme moonlighting in neuronal nuclei. This enzyme, Acss2 (Acetyl-CoA Synthetase 2), is bound to promoters of key neuronal genes in the dorsal hippocampus (a brain region involved in learning and strongly affected by alcohol^30^), and leads to increased histone acetylation and transcription upon alcohol exposure. Strikingly, we showed that this is driven by direct and rapid incorporation of alcohol-derived acetate into histone acetylation, which previously was shown only in the liver^31^. The resulting gene expression changes guide associative memory formation of alcohol-related environmental cues^32^. This is important because drug-associated memories can drive craving and relapse even after protracted periods of abstinence and thus play a central role in driving alcohol consumption and the development of AUD^33-35^. Of note, *ACSS2* polymorphisms were recently linked to AUD in humans^36^.

Importantly, our prior research was limited to passive exposure to ethanol, thus whether this pathway in models of voluntary alcohol drinking that better approximate patterns of human alcohol consumption remains unknown. Therefore, we now assessed the effect of genetic Acss2 inhibition on voluntary alcohol intake and simultaneously assayed the epigenetic and transcriptional changes that accompany voluntary alcohol consumption in key brain regions linked to alcohol use disorder.

## MATERIALS AND METHODS

### Animals

Animal use and all experiments performed were approved by the Institutional Animal Care and Use Committee (IACUC protocols 804849 and 805052). All personnel involved have been adequately trained and are qualified according to the Animal Welfare Act (AWA) and the Public Health Service (PHS) policy. To prevent genetic drift, Acss2 KO (in C57BL/6J genetic background)^37^ and corresponding WT (C57BL/6J strain) controls were generated through homozygous crosses of F1 progeny from heterozygote breeding cages. Mice were housed on a 12h/12h light/dark cycle (7 am to 7 pm), with food and water provided ad libitum. All behavioral experiments were conducted between 7 am and 11 am to reduce time-of-day effects.

### Drinking-in-the-dark paradigm

Single-housed adult mice of both sexes were allowed to voluntarily consume ethanol in the drinking-in-the-dark (DID) paradigm^38^, an extensively used rodent model of binge-like drinking. During the habituation period, mice were acclimated to having two bottles in the housing cage. Following the 1-week habituation period, ethanol intake was evaluated in the DID–2BC (two bottle choice) phase. For the DID–2BC phase, single-housed mice had access to 15% (v/v) ethanol and water for 2 h on days 1-3 and 4 h on day 4. The position of the bottles was alternated daily to avoid side preference. On the second week of DID (DID–1B phase), mice had access to 15% ethanol in a similar schedule, but water was not available during the drinking session (Figure 1A).

**Figure 1.**
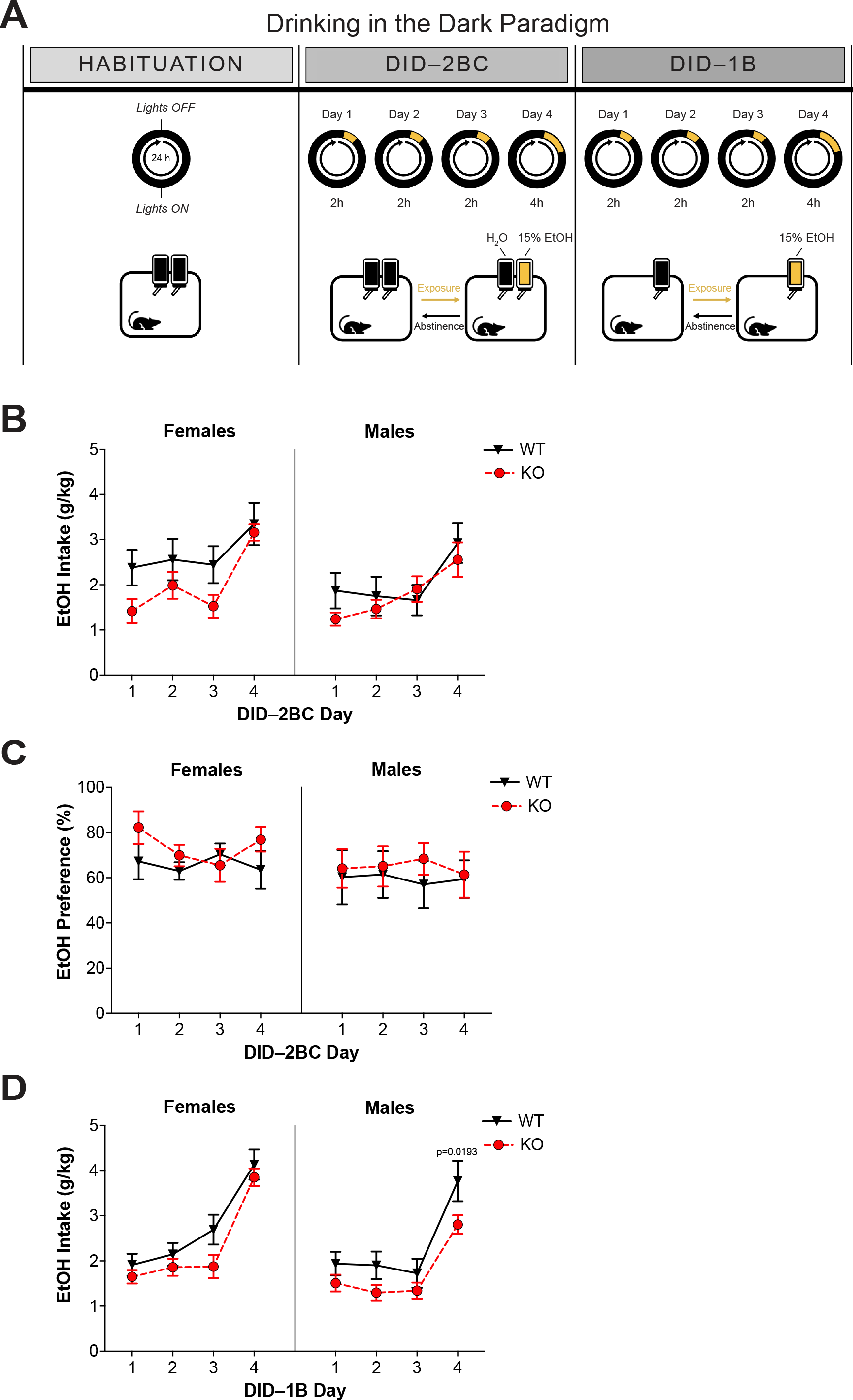
Decreased voluntary alcohol intake in Acss2 KO mice. (A) Schematic depicting the drinking-in-the-dark (DID) behavioral paradigm. Following habituation, mice had access to water and 15% (v/v) ethanol for 4 days (2BC phase), then 15% (v/v) ethanol for 4 days (1B phase). (B) Ethanol intake (g/kg) during the 2B phase in female and male WT and Acss2 KO mice. (C) Ethanol preference (%) in female and male WT and Acss2 KO mice. (D) Ethanol intake (g/kg) during the 1B phase in female and male WT and Acss2 KO mice. During the 1B phase, three-way ANOVA revealed significant effects of both sex (F1,34=4.159, p=0.0493) and genotype (F1,34=11.57, p=0.0017). Post-hoc comparisons showed a significant decrease of alcohol intake on the final day of DID–1B in male mice (p-adj.=0.0193), whereas no statistically significant differences were observed in female mice. For line graphs, symbols and error bars represent the average ± SEM. Sample sizes are as follows: female WT (n=8), female KO (n=10), male WT (n=10), male KO (n=10). 2BC, 2 bottle choice; 1B, 1 bottle; KO, Acss2 KO.

All drinking sessions started 1.0-1.5 h into the dark phase of the light cycle. Bottles were weighed to the nearest hundredth at the beginning and end of each DID session to determine fluid consumption. Mice were weighed prior to each DID session to determine ethanol dose (g/kg). Ethanol consumption was calculated using r15_ethanol=0.97897 g/mL and r100_ethanol=0.789 g/mL. Ethanol preference was calculated as the ratio of ethanol to total fluid intake (mL ethanol solution/mL total fluid).

Animals were sacrificed following the final drinking session via decapitation. Brains were extracted and flash frozen, and subsequently stored at -80C.

### Blood Ethanol Concentration (BEC)

Mice were sacrificed immediately following the final drinking session and blood was collected. Blood ethanol concentration (BEC) was measured using the Ethanol Assay Kit from Sigma-Aldrich® (Cat No. MAK076). Samples were prepared according to manufacturer protocol. Colorimetric readout was done using a BioTek® Synergy™ H1 microplate reader for absorbance at 570 nm.

### RNA-seq

To measure transcriptional responses across different brain regions, we dissected 10-20 mg tissue using a brain matrix and tissue corers from the following brain regions of both male and female WT and Acss2 KO mice: dorsal and ventral hippocampus, dorsal and ventral striatum, amygdala, cortex, and prefrontal (prelimbic/infralimbic) cortex. Total RNA was extracted using TRIzol-chloroform. Total RNA quality was assessed on the Bioanalyzer platform using the RNA 6000 Nano assay (Agilent). mRNA was isolated from 300 ng total RNA using the NEBNext® Poly(A) mRNA Magnetic Isolation Module (E7490L), and libraries were prepared using the NEBNext® UltraTM II RNA Library Prep Kit for Illumina® (E7770).

Sequencing data were aligned to mouse genome assembly GRCm38/mm10 using STAR v2.5.2a with command-line parameters --outFilterType BySJout --outFilterMultimapNmax 20 -- alignSJoverhangMin 8 --alignSJDBoverhangMin 1 --outFilterMismatchNmax 999 -- alignIntronMin 20 --alignIntronMax 1000000 --alignMatesGapMax 1000000 -- readFilesCommand zcat --outFilterMismatchNoverReadLmax 0.04. The STAR index was constructed GENCODE M25 annotation (ENSEMBL 100 assembly). Data were filtered for poor mapping tags using samtools v1.1 subcommand view `-bS -q 255`. Technical repeat samples from two flowcells were combined using samtools merge and aligned tags were sorted by chromosome position and indexed using samtools sort and index, respectively. HTseq v0.6.1 was used to quantify tag counts over exons with command-line parameters `-f bam -r pos -s reverse -t exon -i gene_id --additional-attr=gene_name`. Differentially expressed genes (DEGs) between ACSS2 KO and WT samples were detected separately in each tissue using DESeq2^39^ in R v4.0.2 with the FDR controlled at 0.1.

Heatmaps were generated using R library pheatmap and PCA was rendered from the full data matrix with zero rows removed, using ggplot2. Volcano plots were also generated using ggplot2. Gene Ontology analysis was performed using DAVID^40,41^ with the FDR controlled at 0.1. RNA-seq data were visualized on the UCSC Genome Browser^42^ (mm10, GRCm38 assembly); tracks were created using DeepTools^43^ bamCoverage v3.4.3 with command-line parameter `--scaleFactor` set to the reciprocal DESeq2 size factor for that sample. Finally, bigWigs from biological repeats were combined using wiggletools (“mean” function) and then rendered from bedGraph to bigWig using UCSC Genome Browser tools’ bedGraphToBigWig.

### H3K27ac ChIP-seq

Approximately 20 mg of ventral striatal tissue from each mouse was minced on ice, cross-linked with 1% formaldehyde for 10 min and quenched with 125 mM glycine for 5 min. Nuclei were prepared by dounce homogenization of cross-linked tissue in nuclei isolation buffer (50 mM Tris-HCl at pH 7.5, 25 mM KCl, 5 mM MgCl2, 0.25 M sucrose) with freshly added protease inhibitors and sodium butyrate. Nuclei were lysed in nuclei lysis buffer (10 mM Tris-HCl at pH 8.0, 100 mM NaCl, 1 mM EDTA, 0.5 mM EGTA, 0.1% sodium deoxycholate, 0.5% *N*-lauroylsarcosine) with freshly added protease inhibitors and sodium butyrate, and chromatin was sheared to approximately 250 bp in size using a Covaris S220 sonicator. Equal aliquots of sonicated chromatin were used per immunoprecipitation reaction with 4 μl H3K27ac antibody (Abcam; 4729) preconjugated to Protein G Dynabeads (Life Technologies). Ten percent of the chromatin was saved as input DNA. ChIP reactions were incubated overnight at 4°C with rotation and washed three times in wash buffer. Immunoprecipitated DNA was eluted from the beads, reversed cross-linked and purified together with the input DNA. Exactly 10 ng DNA (either ChIP or input) was used to construct sequencing libraries using the NEBNext Ultra II DNA library preparation kit for Illumina (New England Biolabs; #E7760). Libraries were multiplexed using NEBNext Multiplex Oligos for Illumina (dual index primers) and paired-end sequenced (42 bp) on the NextSeq 500 platform (Illumina) in accordance with the manufacturer’s protocol.

Sequencing data were aligned to mouse genome assembly GRCm38/mm10 using bowtie2^44^ v2.3.4.3 with command-line parameter `--local`. Data were filtered for poor mapping tags using samtools^45^ v1.1 subcommand view `-bS -q 5 -f 2`. Data were sorted by tag name using samtools sort -n and PCR duplicates were removed using PICARD MarkDuplicates v 2.21.3-SNAPSHOT with command-line parameters `REMOVE_DUPLICATES=True ASSUME_SORT_ORDER=queryname`. Aligned, deduplicated tags were sorted by chromosome position and indexed using samtools sort and index, respectively. RGT-THOR^46^ v0.13.2 was used to detect differentially enriched peaks using input samples to control for sonication efficiency artifacts at regions of open chromatin. RPM-adjusted counts over each differential peak were then assessed using a python v3 script. Peaks were associated to the nearest gene using HOMER^47^ v4.6 annotatePeaks.pl and reported annotation was used to construct genomic distribution pie charts in Supplementary Figure 3A-B.

Peaks were filtered for a 4x alteration in ACSS2 KO vs WT and the losses were scanned for motif enrichment using HOMER findMotifsGenome.pl using command-line parameters `-size 300 -mask` and using the 4x gained peaks as background. The reverse scan was also carried out for the gained peaks, using the losses as background. ChIP-seq data were visualized using UCSC Genome Browser; tracks were created using DeepTools bamCompare v3.4.3 with command-line parameters `-bs 1 --effectiveGenomeSize 2652783500` and setting input to the background using parameter `-b2`. Gene Ontology analysis was performed using DAVID with the FDR controlled at 0.1. Boxed violin plots relating H3K27ac changes to gene expression changes were created using the ggplot2 library in R v4.0.2. Heatmaps of lost or gained H3K27ac peaks reflect a 2kb window around the peak center and were created using python v3 scripts (vector quantitation and visualization using the PIL library).

## RESULTS

### Loss of Acss2 reduces voluntary alcohol intake, especially in males

To investigate voluntary alcohol intake, we employed the “drinking in the dark” (DID) paradigm (Figure 1A), which is widely used to model binge drinking in mice. Here, animals typically consume enough alcohol to reach high blood ethanol concentrations (BECs) and exhibit behavioral manifestations of intoxication^38^. We used a modified version of DID in which mice had access to 15% (v/v) ethanol and water during days 1-4 (two bottle choice, 2BC phase), and only 15% (v/v) ethanol during days 5-8 (one bottle, 1B phase) (Figure 1A). We tested both female and male mice to uncover potential sex-specific effects. To investigate the role of ACSS2 during drinking behavior, we used a constitutive Acss2 knockout (KO) line we recently developed on C57BL6/J background^37^. Importantly, Acss2 knockout is well tolerated and the transgenic mouse exhibits no overt behavioral abnormalities or transcriptional and epigenetic changes at baseline.

Acss2 KO had no effect on ethanol intake or preference during the 2BC phase of the DID model (Figure 1B,C). During the 1B phase, however, we found that Acss2 KO decreased alcohol drinking in a sex-specific manner (Figure 1D). Three-way ANOVA revealed significant effects of both sex (F_1,34_=4.159, p=0.0493) and genotype (F_1,34_=11.57, p=0.0017): Acss2 KO mice consumed less alcohol compared to WT littermates, and this effect was primarily driven by males. Post-hoc comparisons showed a significant decrease of alcohol intake on the final day of DID in male mice (Figure 1D, p-adj.=0.0193), while no statistically significant differences were observed in female mice.

Mice were sacrificed immediately following the final 1B session and blood was collected to measure BEC. Intriguingly, we found that deletion of Acss2 had a sex-specific effect: While no difference was observed in male mice, female Acss2 KO mice had significantly higher BECs (student’s t-test, p=0.0287) compared to WT littermates (Supplementary Figure 1A). Taken together with the alcohol intake data (Figure 1), the observed elevated BEC in female mice is potentially due to inhibition of liver alcohol metabolism^48^ in the Acss2 KO animals.

### Blunted alcohol drinking-related transcriptional states in the ventral striatum of Acss2 KO mice compared to WT

Next, we performed RNA-seq to profile transcriptional changes that accompany voluntary alcohol consumption across various key brain regions linked to AUD. We sacrificed mice following the final, 4-hour drinking session and queried gene expression in male and female WT and Acss2 KO mice across the brain, focusing on regions implicated in alcohol use disorder^49^. These included the ventral striatum (which mediates alcohol reward), dorsal striatum (linked to compulsive drinking), ventral hippocampus (mediating stress, emotion and affect), dorsal hippocampus (which confers alcohol-related learning), cortex (linked to executive control), and amygdala (implicated in withdrawal-related negative affective outcomes). As expected, our principal component analysis revealed marked variance of gene expression across these brain regions investigated in the Acss2 KO in both male and female animals (Supplementary Figure 2A,B). In all brain regions and in both sexes, *Acss2* was the most significantly downregulated gene, with clear average loss of expression across all tissues (males shown in Figure 2A,B and Supplementary Figure 2E, females shown in Supplementary Figure C).

**Figure 2.**
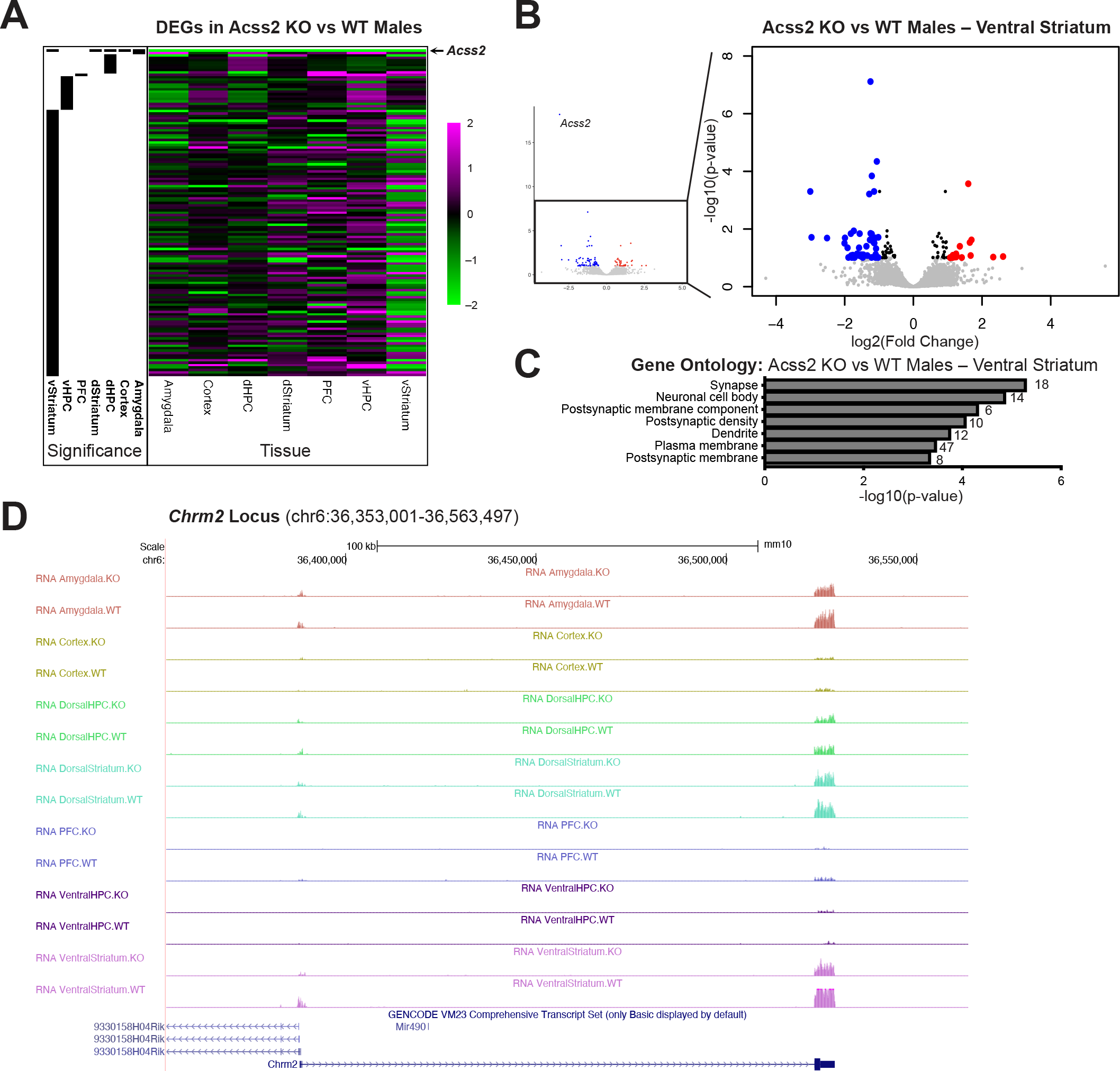
Transcriptional profiling across key brain regions following voluntary alcohol intake in male Acss2 KO and WT mice. (A) Heatmap showing differentially expressed genes (DEGs, n=135) across seven brain regions investigated in Acss2 KO vs WT male mice following voluntary alcohol intake. Black bars (left) denote the region corresponding to significant DEGs (FDR<0.1). For the heatmap, magenta (2) indicates increased expression in Acss2 KO vs WT males, black (0) indicates no differential expression between Acss2 KO vs WT males, and green (-2) indicates decreased expression in Acss2 KO vs WT males. (B) Volcano plot of ventral striatal DEGs (n=111, with (thumbnail) and without (zoom-in) *Acss2*) identified between Acss2 KO vs WT males. Each dot represents a gene. Red dots denote significantly upregulated genes, and blue dots denote significantly downregulated genes. (C) Gene Ontology analysis of ventral striatal DEGs between Acss2 KO vs WT males showing enrichment of genes linked to synaptic functions. The top 7 most significant GO terms are displayed. (D) Gene expression tracks of the *Chrm2* locus (chr6:36,353,001-36,563,497) across brain regions in male mice, showing specific significant decrease of *Chrm2* expression in the ventral striatum. DEGs, differentially expressed genes; dHPC, dorsal hippocampus; dStriatum, dorsal striatum; PFC, prefrontal cortex; vHPC, ventral hippocampus; vStriatum, ventral striatum; GO, Gene Ontology.

In males, we found the highest number of differentially expressed genes (DEGs, n=111) in the ventral striatum (Figure 2A and Supplementary Table 1). Other brain regions showed considerably fewer DEGs between Acss2 KO and WT: 14 DEGs in the ventral hippocampus, 9 DEGs in the dorsal hippocampus, 2 DEGs in the amygdala, and 1 DEG each (*Acss2*) in the dorsal striatum, cortex and prefrontal cortex (Supplementary Table 1). In the ventral striatum, the majority of DEGs (n=71) were significantly downregulated in the Acss2 KO animals compared to WT littermates (Figure 2B). Intriguingly, several of the ventral striatal DEGs were previously identified in human AUD patients, including downregulated *Akap12* (*A-kinase anchoring protein 12*), as well as upregulated *Camkk2* (*Ca/calmodulin-dependent protein kinase kinase 2*) and *Mef2C* (*myocyte enhancer factor 2C*)^50^. Further, we found a significant enrichment of delayed primary response genes and secondary response genes^51^ among the ventral striatal DEGs (hypergeometric p=0.000148). These included *Hcrtr2* (*hypocretin receptor 2*; Supplementary Figure 3A), *Nap1l5* (*nucleosome assembly protein 1-like 5*; Supplementary Figure 3B) and *Stac* (*SH3 and cysteine rich domain*; Supplementary Figure 3C). The downregulation of these genes suggests blunting in the Acss2 KO of alcohol-induced transcriptional remodeling in the ventral striatum, which might contribute to the observed differences in drinking behavior (Figure 1). This was further supported by Gene Ontology (GO) analysis, which revealed that downregulated ventral striatal DEGs were linked to synaptic functions, postsynaptic components, and postsynaptic density/membrane (Figure 2C). For example, we found decreased expression in Acss2 KO of delayed primary response gene *Chrm2*, a gene encoding for synaptic muscarinic acetylcholine receptor M2 (Figure 2D). Chrm2 is strongly implicated in memory and cognitive function, and mutations of this gene were shown to predispose carriers to alcohol dependence and affective disorders^52^.

By contrast, transcriptional changes between WT and Acss2 KO were more limited in female mice. This was in line with the less pronounced behavioral differences observed in females (Figure 1). We found 69 DEGs in the prefrontal cortex, 12 DEGs in the dorsal hippocampus, 3 DEGs in the cortex, 2 DEGS each in the amygdala and ventral hippocampus, and 1 DEG each (*Acss2*) in the dorsal striatum and ventral striatum (Supplementary Table 1). The prefrontal cortical DEGs were enriched for genes related to myelination (Supplementary Figure 2D), and only one was a delayed response gene (*Nrn1*). Considering all brain regions, only 9 genes were differentially expressed in both males and females (Supplementary Table 1). All of these have been previously implicated in AUD (see discussion).

### Depleted H3K27ac correlates with blunted gene expression programs in the ventral striatum of Acss2 KO mice following voluntary alcohol intake

Previously, we showed that Acss2 drives alcohol-related histone acetylation in the dorsal hippocampus, leading to gene expression changes that underlie alcohol-associated learning^32^. Whether Acss2-dependent histone acetylation plays a similar role in other brain regions that mediate other critical aspects of drinking behavior, however, remains unknown.

To test this possibility in the ventral striatum, we performed chromatin immunoprecipitation coupled with high throughput sequencing (ChIP-seq) targeting H3K27ac, a histone acetylation mark we previously showed to be induced by alcohol-derived acetate^32^. We focused on male mice given more pronounced behavioral (Figure 1) and transcriptional (Figure 2) dysregulation described above. Comparing H3K27ac enrichment in the ventral striatum of male WT and Acss2 KO mice following DID, we found that H3K27 acetylation was markedly decreased in Acss2 KO animals (Figure 3A): we found 8,061 significantly depleted peaks in Acss2 KO mice compared to WT mice, and only 4,298 significantly enriched peaks (passing genome-wide statistical significance and exhibiting at least 4x fold change). We found a similar genomic distribution of depleted (Supplementary Figure 4A) and enriched peaks (Supplementary Figure 4B), with a slightly higher proportion of depleted peaks in intergenic space. Gene Ontology (GO) analysis of genes near altered peaks revealed enrichment of genes related to neuronal axonogenesis near reduced H3K27ac peaks, and enrichment of genes related to synaptic membrane adhesion associated with increased H3K27ac peaks (Supplementary Figure 4C-D).

**Figure 3.**
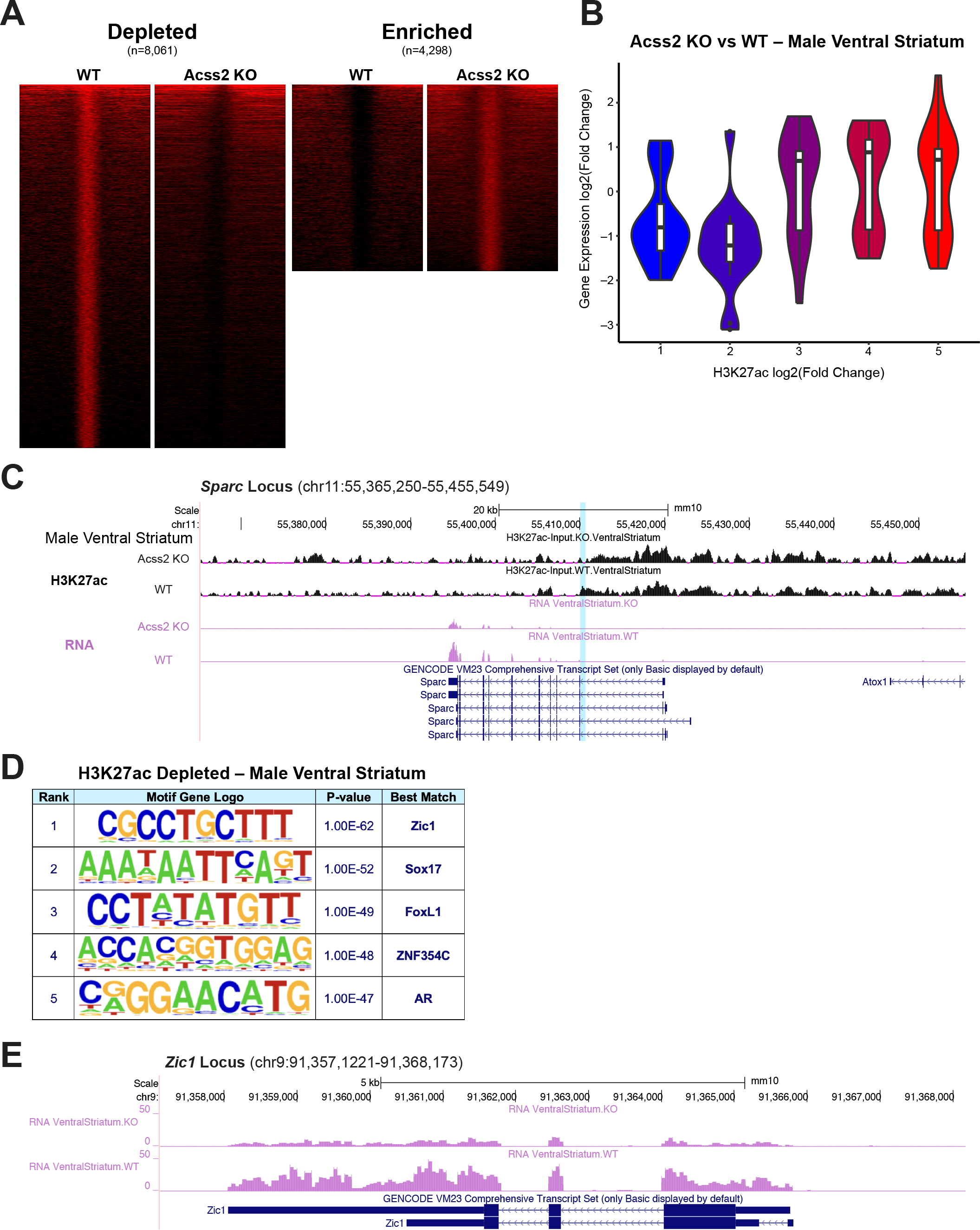
H3K27ac profiling of the ventral striatum following voluntary alcohol intake in male WT and Acss2 KO mice. (A) Heatmaps showing 2 kb windows of significantly depleted (n=8,061) and significantly enriched (n=4,298) H3K27ac peaks in the ventral striatum of Acss2 KO vs WT male mice following voluntary alcohol consumption. (B) Violin plots showing positive correlation between H3K27ac enrichment changes (ChIP-seq, x-axis) and gene expression changes (RNA-seq, y-axis) in the ventral striatum of Acss2 KO vs WT male mice following voluntary alcohol consumption. (C) Tracks of the *Sparc* locus (chr11:55,365,250-55,455,549) showing decreased H3K27ac enrichment (top two rows) and decreased gene expression (bottom two rows) in the Acss2 KO male ventral striatum compared to that of WT. (D) *De novo* motif analysis showing the top 5 significantly enriched transcription factor binding motifs in H3K27ac depleted peaks. (E) Gene expression tracks of the *Zic1* locus (chr9:91,357,1221-91,368,173).

We next integrated our transcriptional and epigenetic datasets by comparing male ventral striatal gene expression with H3K27ac enrichment (Supplementary Figure 4E-G). As expected, genes with lower H3K27ac tended to show decreased transcription (Figure 3B, with H3K27ac binning shown in Supplementary Figure 4F), suggesting that the blunted drinking-related transcriptional states described above (Figure 2) are related to the depletion of H3K27ac in the ventral striatum of Acss2 KO male mice following voluntary ethanol intake. For example, we found decreased H3K27ac at a putative enhancer flagged in ENCODE (Figure 3C, top two tracks, blue highlight) of the *Sparc* gene (*Secreted protein acidic rich in cysteine*), the expression of which was a significantly decreased (Figure 3C, bottom two tracks). Notably, *Sparc* was previously implicated in models of chronic intermittent alcohol exposure in the prefrontal cortex^53^.

We then performed *de novo* motif analysis of both lost and gained H3K27ac peaks in the ventral striatum, and found that the binding motif of transcription factor Zic1 (zinc finger protein of the cerebellum 1) was the most enriched in H3K27ac peaks that were depleted in Acss2 KO animals (Figure 3D). Of note, the expression of *Zic1* itself was significantly decreased in the male ventral striatum but not in other brain regions (Figure 3E), suggesting that this transcription factor might act as a regulator of alcohol-driven histone acetylation in the ventral striatum. Interestingly, no differences of *Zic1* expression were observed in female mice (Supplementary Table 1).

## DISCUSSION

Here, we show that mice with knockout of Acss2 consume significantly less alcohol in models of voluntary drinking. By performing transcriptional profiling across AUD-associated brain regions, we find that decreased drinking is associated with blunted gene expression in the ventral striatum, a brain region closely linked to alcohol and drug reward. We find that both behavioral and transcriptional differences between WT and Acss2 KO mice are more pronounced in male compared to female mice following voluntary alcohol intake. Further, we identify decreased acetylation of histone H3 lysine 27 (H3K27) as a potential epigenetic mechanism in the ventral striatum of male Acss2 KO mice.

Intriguingly, we found significantly decreased alcohol intake during the one bottle (1B) phase but not during the two-bottle choice (2BC) phase of the drinking-in-the-dark paradigm. This is surprising, as alcohol drinking phenotypes generally show noticeable correlation between the different phases of DID measuring alcohol intake and alcohol preference^54^. However, in our experiments, the 2BC phase always preceded the 1B phase, it is possible that the different behavioral outcomes were due to differences in alcohol drinking history.

Whether longer exposure to 2BC or switching the order of the 1B and 2BC phases would manifest in decreased alcohol preference and decreased drinking during the 2BC phase in Acss2 KO mice, remains to be seen.

Consistent with the literature, we observed slightly elevated levels of alcohol intake in female compared to male mice (Figure 1B,D, Supplementary Figure 1). While both sexes escalate alcohol intake during DID, females generally tend to drink more and achieve higher blood ethanol concentrations^38^. Intriguingly, the phenotypic differences between Acss2 KO mice and WT littermates were more pronounced in male compared to female mice in our experiment (Figure 1B-D). This was accompanied by more severe transcriptional dysregulation, especially in the ventral striatum, suggesting that Acss2 plays an important role in mediating alcohol reward and consumption, especially in males. Surprisingly, we found that female Acss2 KO mice had significantly elevated blood ethanol concentrations (BEC) at the end of the DID paradigm compared to WT, a difference we did not observe in males (Supplementary Figure 1). The significantly elevated BEC in female mice is potentially due to feedback inhibition of alcohol metabolism in the Acss2 KO animals^48^. These mice lack Acss2 expression across all tissues including the liver, which could result in slower breakdown of alcohol due increased levels of acetate or acetaldehyde, thus resulting in elevated BEC. As male Acss2 KO mice drank significantly less alcohol, this effect on BEC was only observed in females.

In line with the more subtle behavioral effects, we found less pronounced transcriptional dysregulation in the brain of female Acss2 KO mice. Intriguingly, while only 9 DEGs overlapped between male and female mice across all tissues, all of these genes have been previously implicated in AUD. In addition to *Acss2*^36^, we found changes in the expression of *Trf* (transferrin; carbohydrate deficient transferrin has been proposed as a biomarker of alcohol use^55^) and *Atp1a1*, which is increased following prenatal exposure to alcohol^56^. Strikingly, six of the nine shared DEGs between males and females were strongly linked to myelinization. *Mbp* (myelin basic protein), *Mag* (myelin-associated glycoprotein) and *Plp* (proteolipid protein) are decreased in the brain of human AUD patients^57^. *Mal* (myelin and lymphocyte protein) expression is decreased in mice following binge drinking^58^, and expression of *Kcnv1* (potassium voltage-gated channel modifier subfamily V member 1) correlates with voluntary ethanol consumption in both the cortex and the ventral striatum^59^. *Mobp* (myelin-associated oligodendrocyte basic protein) is decreased both in humans and animal models of alcohol use^60^. *Acss2* itself is heavily implicated in lipid metabolism, both as a transcriptional regulator and as an important source of intracellular acetyl-CoA^61^. While we have previously shown that this enzyme is primarily nuclear localized in differentiated neurons^62^, it remains to be seen whether potential effects on cytoplasmic metabolism, lipid synthesis and myelinization also contribute to the behavioral differences observed in Acss2 KO mice, especially in the context of alcohol consumption.

An interesting category of genes enriched among the ventral striatal DEGs in males were delayed primary response genes or secondary response genes^51^. These are distinct groups of activity-regulated genes in neurons, which are induced in response to neuronal activation and have been linked to transcription-dependent synaptic plasticity, long-term potentiation, long-term depression, and synaptic scaling^63-65^. Notably, several of the DEGs we identified in these categories have been previously implicated in alcohol use. For example, in humans, the *CHRM2* gene predisposes to alcohol dependence, drug dependence^52^, as well as affective disorders including major depressive disorder^66^. *Hcrtr2* encodes for the receptor of neuropeptide hypocretin/orexin, which is known to be dysregulated in alcohol dependence^67^ and compulsive alcohol drinking in mice^68^. Further, Hcrt signaling in the amygdala was shown to be necessary for alcohol-seeking behavior in mouse models of alcohol dependence^68^.

In line with blunted gene expression programs, we found that acetylation of H3K27 tended to be depleted in the ventral striatum of male mice following voluntary alcohol intake. We have previously shown that Acss2 mediates alcohol-driven histone acetylation in the brain by converting alcohol-derived acetate into acetyl-CoA, which is deposited on histones by histone acetyltransferases^32^. Importantly, our current observations of decreased H3K27ac in Acss2 KO animals align with these findings. Our *de novo* motif analysis of depleted H3K27ac peaks identified the most enriched binding site being that of transcription factor Zic1. Interestingly, mRNA levels of *Zic1* itself were significantly decreased in the KO mice. Interestingly, Zic1 has been previously implicated in brain development^69^ and Zic1-deficient mice show hypoplastic cerebellum and spinal cord^70^. Here, we identify a potential new role of Zic1 in the ventral striatum to mediate alcohol-related transcriptional adaptations that could underlie voluntary drinking. In future experiments, we will test whether inhibition or loss of Zic1 phenocopies Acss2 loss during drinking-in-the-dark or similar behavioral assays.

Taken together, our findings indicate that knockout of Acss2 results in decreased voluntary alcohol intake, especially in male mice. We show that this behavioral phenotype is related to blunted alcohol-related gene expression in the ventral striatum, driven by diminished histone acetylation in this brain region. Our findings further emphasize the critical role of Acss2 in alcohol metabolism and related epigenetic, transcriptional, and behavioral changes, which warrant further exploration of this pathway as a potential future therapeutic target in individuals struggling with AUD.

## FIGURE LEGENDS

**Supplementary Figure 1.**
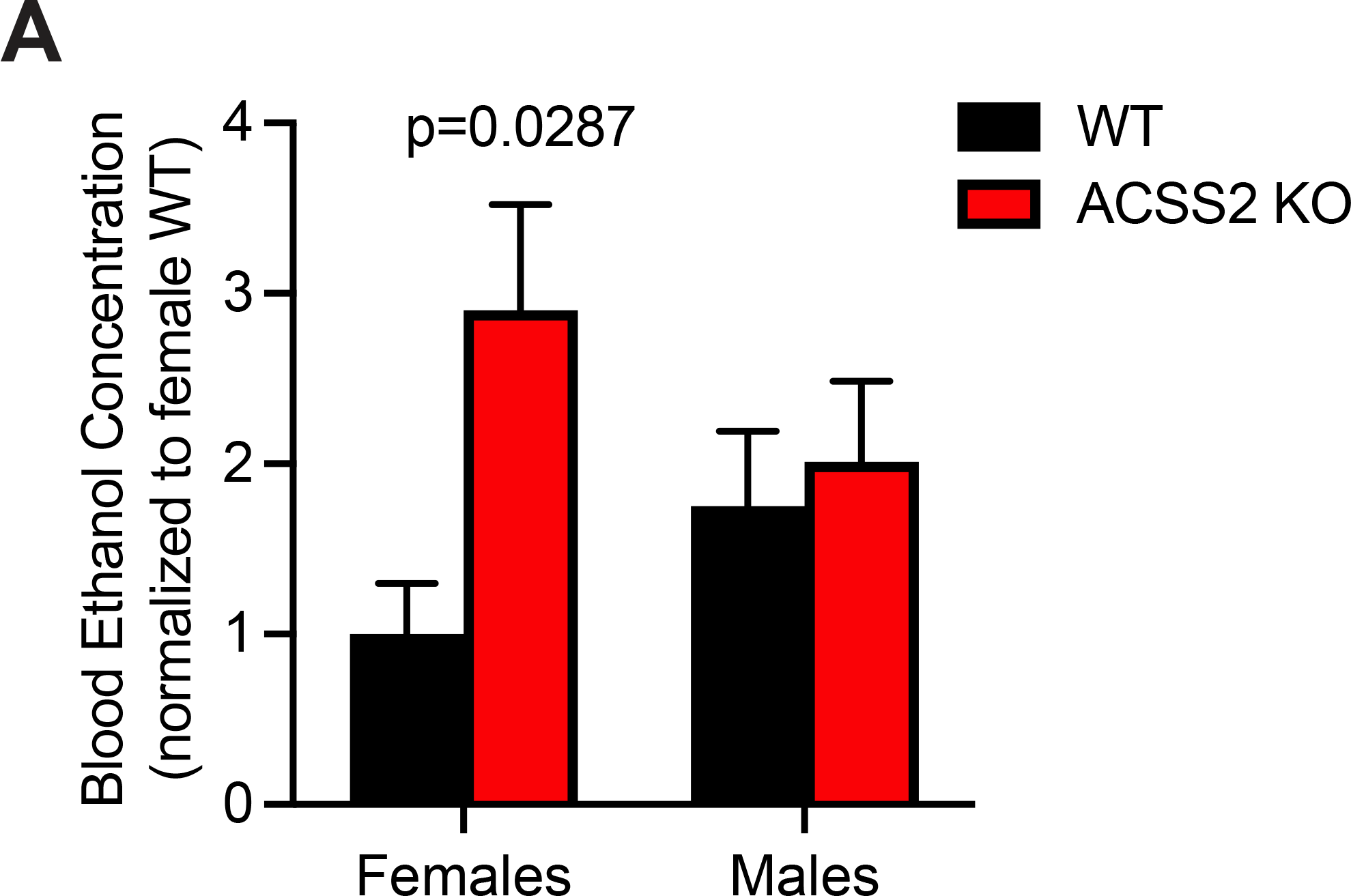
Blood ethanol concentrations following the final session of the drinking-in-the-dark paradigm. (A) Average blood ethanol concentration (BEC; normalized to WT females) following the final session of the DID–1B phase in female and male WT and Acss2 KO mice. Female Acss2 KO animals had significantly higher BEC on average compared to female WT littermates (student’s t-test, p=0.0287), whereas no statistically significant differences were observed in male mice. Columns and error bars represent the average ± SEM. Sample sizes are as follows: female WT (n=7), female KO (n=9), male WT (n=10), male KO (n=10). BEC, blood ethanol concentration; DID, drinking in the dark; KO, Acss2 KO.

**Supplementary Figure 2.**
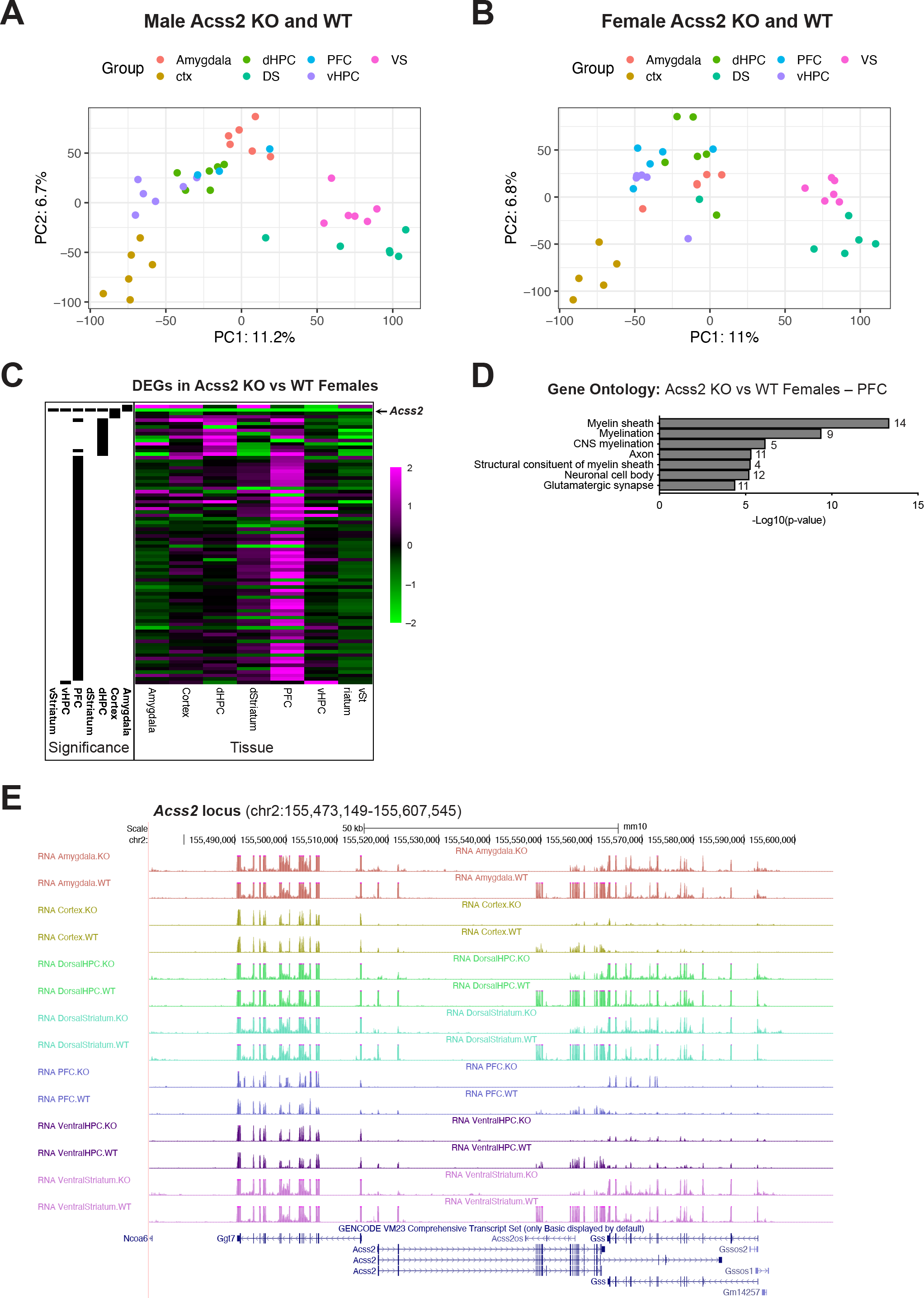
Transcriptional profiling of key brain regions following voluntary alcohol intake in Acss2 KO and WT mice. (A) Principal component analysis of male gene expression data showing clear transcriptional separation across various brain regions investigated. (B) Principal component analysis of female gene expression data showing clear transcriptional separation across various brain regions investigated. (C) Heatmap showing differentially expressed genes (DEGs, n=82) across seven brain regions investigated in Acss2 KO vs WT female mice following voluntary alcohol consumption. Black bars (left) denote significant DEGs corresponding to each region (FDR<0.1). Magenta (2) indicates increased expression in Acss2 KO vs WT females, black (0) indicates no differential expression between Acss2 KO vs WT females, and green (-2) indicates decreased expression in Acss2 KO vs WT females. (D) Gene Ontology analysis of prefrontal cortical DEGs between Acss2 KO vs WT females showing enrichment of genes linked to myelination. (E) Gene expression tracks showing reproducible loss of *Acss2* expression (chr2:155,473,149-155,607,545) across all brain regions in male Acss2 KO compared to WT mice. Ctx, cortex; dHPC, dorsal hippocampus; DS or dStriatum, dorsal striatum; PFC, prefrontal cortex; vHPC, ventral hippocampus; VS or vStriatum, ventral striatum; PC, principal component; DEGs, differentially expressed genes.

**Supplementary Figure 3.**
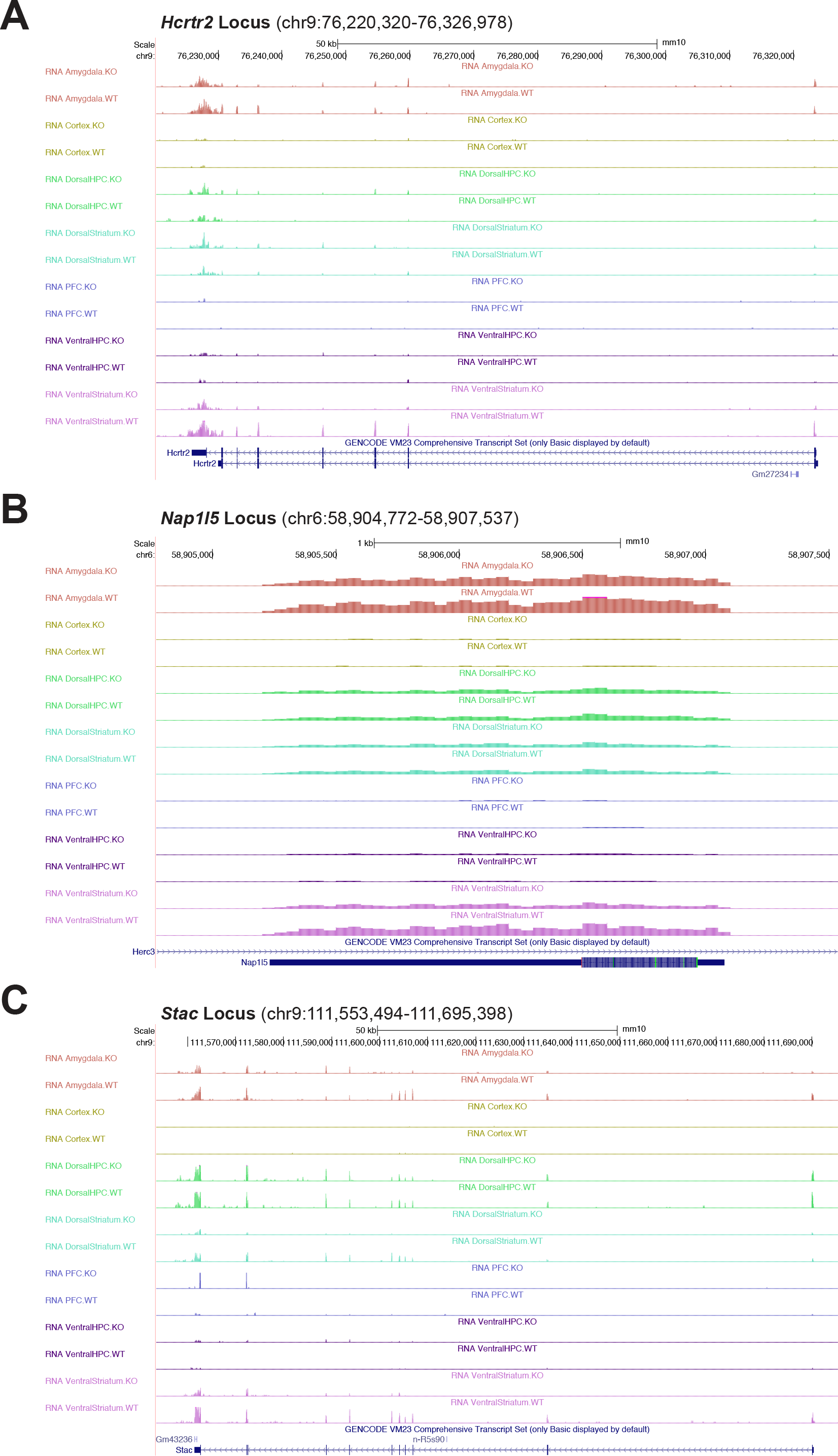
Delayed response genes are expressed at significantly lower levels in the ventral striatum of male Acss2 KO mice following voluntary alcohol intake. (A) Gene expression tracks showing significantly decreased expression of known delayed response genes (A) *Hcrtr2* (chr9:76,220,320-76,326,978), (B) *Nap1I5* (chr6:58,904,772-58,907,537), and (C) *Stac* (chr9:111,553,494-111,695,398) in the ventral striatum of Acss2 KO and WT mice following voluntary alcohol intake.

**Supplementary Figure 4.**
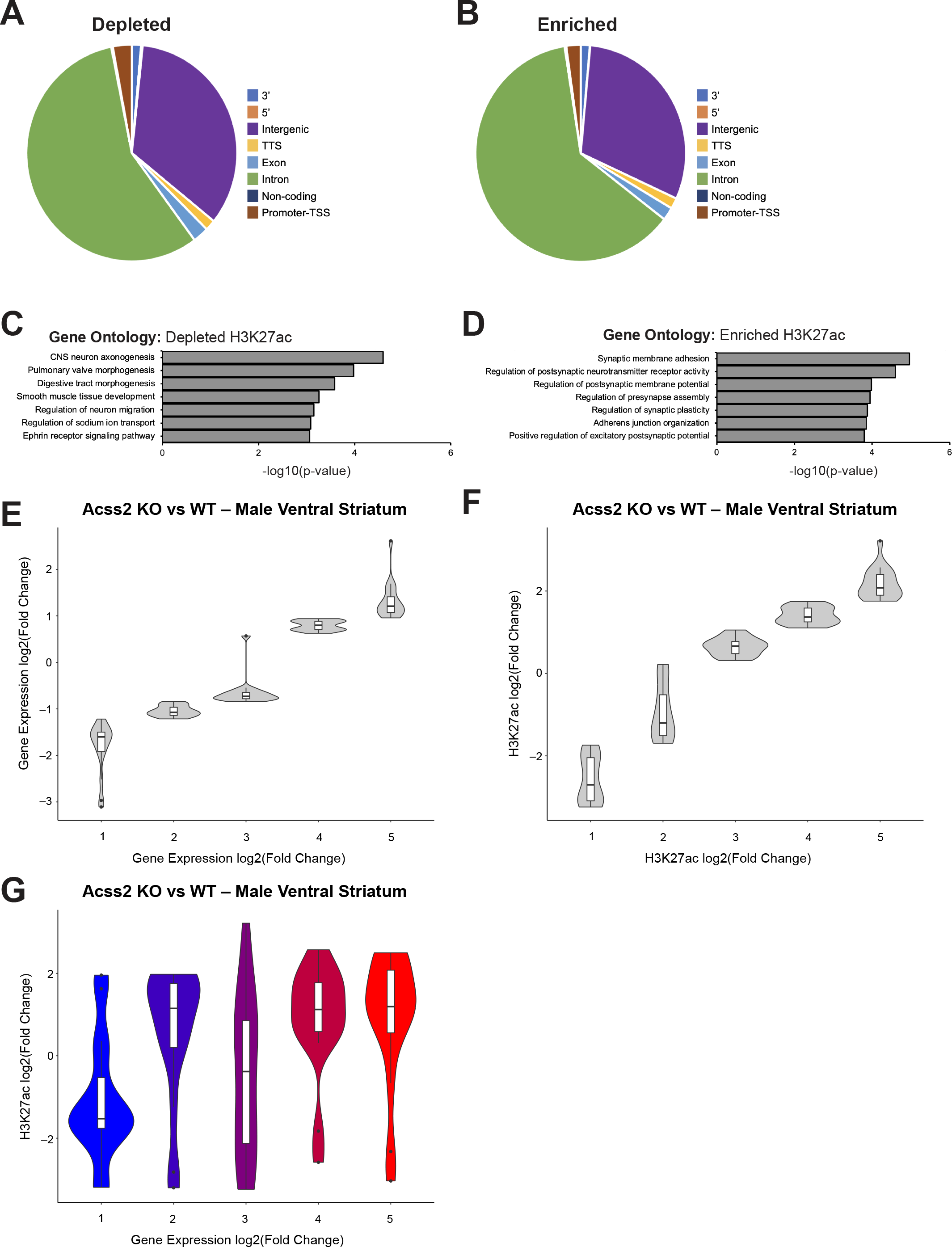
Epigenetic profiling of the ventral striatum following voluntary alcohol intake in male Acss2 KO and WT mice. (A,B) Pie charts showing the genomic distribution of (A) significantly depleted and (B) significantly enriched H3K27ac peaks in the ventral striatum of male Acss2 KO mice following voluntary alcohol consumption. Peaks showing genome-wide significance and at least 4x fold change were considered significant. (C,D) Gene Ontology analysis of (C) depleted and (D) enriched H3K27ac peaks in the ventral striatum of Acss2 KO mice following voluntary alcohol intake. (E-G) Construction plots showing quintiles of (E) gene expression, (F) H3K27ac enrichment, and (G) H3K27ac enrichment (ChIP-seq, y-axis) vs gene expression (RNA-seq, x-axis) in the ventral striatum of male Acss2 KO vs WT mice following voluntary alcohol intake.

**Supplementary Table 1.**List of differentially expressed genes between WT and Acss2 KO following voluntary alcohol intake in male and female mice across brain regions. Ctx, cortex; dHPC, dorsal hippocampus; DS, dorsal striatum; PFC, prefrontal cortex; VS, ventral striatum; LFC, log-fold change.

## DATA AVAILABILITY

Sequencing data are publicly available on Gene Expression Omnibus under GSE249119.

